# Altitudin S from *Bacillus altitudinis* ECC22 defines a new subgroup of circular bacteriocins

**DOI:** 10.1101/2025.10.31.685828

**Authors:** Ester Sevillano, Irene Lafuente, Nuria Peña, Luis M. Cintas, Estefanía Muñoz-Atienza, Pablo E. Hernández, Juan Borrero

## Abstract

Bacteriocins are ribosomally synthesized antimicrobial peptides exhibiting diverse structures and mechanisms of action. *Bacillus altitudinis ECC22*, previously shown to produce the circular bacteriocins pumilarin and altitudin A, was found to harbor an additional biosynthetic gene cluster encoding a novel circular bacteriocin, designated altitudin S Proteomic analysis of active supernatant fractions confirmed the production of altitudin S, with a molecular mass of 8379 Da, consistent with head-to-tail cyclization. The peptide is synthesized as a 132-residue precursor comprising a 56-amino-acid leader and a 76-residue circular mature core. Structural modeling predicted a compact saposin-like fold composed of five α-helices and a strongly cationic surface (pI ≈ 11.0, net charge +13). Altitudin S was successfully produced using a cell-free protein synthesis system coupled to split-intein mediated ligation (IV-CFPS/SIML) and exhibited a narrow but reproducible antimicrobial spectrum. Comprehensive sequence, structural, and phylogenetic analyses revealed that altitudin S is a highly divergent circular bacteriocin, defined by distinctive sequence features and physicochemical properties, including an exceptionally high isoelectric point, net charge, and low hydrophobicity. Bioprospecting across sequence databases identified homologs of altitudin S in diverse *Bacillales* species, all showing high sequence similarity, conserved structural features and preservation of its distinctive physicochemical profile. Genomic analysis further revealed a conserved biosynthetic gene cluster among all altitudin S homologs, notably including a gene encoding a characteristic M48-family metallopeptidase. Altogether, these findings support the classification of altitudin S and its homologs as representatives of a novel subgroup of circular bacteriocins.

## 1. Introduction

The increasing prevalence of antimicrobial resistance (AMR) constitutes a major global health crisis, undermining the efficacy of current antibiotics and severely restricting therapeutic options for bacterial infections [1]. Multidrug-resistant pathogens have emerged across clinical, agricultural, and environmental settings, further intensifying the urgent demand for novel antimicrobial strategies [2]. Despite this need, the discovery of new antibiotics has markedly declined in recent decades, hindered by scientific, economic, and regulatory challenges. Consequently, alternative approaches are gaining increasing attention, with bacteriocins standing out as promising candidates to combat resistant bacteria, either as standalone therapeutics or in synergistic combination with conventional antibiotics [3].

Bacteriocins are ribosomally synthesized antimicrobial peptides produced by bacteria to inhibit the growth of closely related or competing bacterial species. They exhibit remarkable structural diversity and a wide range of mechanisms of action, making them attractive candidates for diverse biotechnological and clinical applications. Bacteriocins are classified into major groups, with class I bacteriocins—posttranslationally modified peptides— being of particular interest due to their structural complexity and functional diversity. Within this class, circular bacteriocins form a distinct subclass characterized by a head-to-tail cyclized backbone, a unique feature that confers exceptional stability, protease resistance, and tolerance to broad pH ranges and elevated temperatures [4,5]. Circular bacteriocins are typically categorized into two main subgroups based on their sequence features and physicochemical properties. Subgroup I members typically consist of mature peptides of 60–70 amino acids with variable leader sequences (2–49 amino acids). They exhibit high isoelectric points (pI 9.5–10.5), net positive charges of +2 and +6, and high aliphatic indices (>115), together with moderately positive GRAVY values (0.5–1.1). These properties support strong electrostatic interactions with negatively charged bacterial membranes, as well as enhanced membrane insertion and disruption due to their hydrophobicity. In contrast, subgroup II circular bacteriocins are shorter, with mature cores of about 58 amino acids and leader peptides of 22–35 residues. They display broader variability in pI (4.0–10.0) and include peptides with neutral or even negative net charges, suggesting alternative modes of action and target specificity. This diversity in charge and hydrophobicity reflects the evolutionary and functional heterogeneity within this group of antimicrobial peptides [6]. Beyond their structural and physicochemical diversity, circular bacteriocins are typically encoded within specific genetic clusters that contain the essential genes for their synthesis, transport, and immunity. A conserved hallmark of these clusters is the presence of the stage II sporulation protein M, a member of the DUF95 superfamily, which is thought to play a crucial role in their biosynthesis and cyclization [7]. Functionally, circular bacteriocins have been identified in multiple bacterial genera and display potent antimicrobial activity against Gram-positive bacteria, including antibiotic-resistant pathogens [8].

In this study, we report the characterization of a novel circular bacteriocin, altitudin S, produced by *Bacillus altitudinis* ECC22, a strain previously shown to synthetize two other circular bacteriocins: pumilarin and altitudin A [9]. An integrated strategy combining genome-based bioinformatic analysis, chromatographic purification, and mass spectrometry confirmed the production and circular conformation of altitudin S by *B. altitudinis* ECC22. Altitudin S was further synthesized using an in vitro cell-free protein synthesis (IV-CFPS) system coupled to a split intein mediated ligation (SIML) [10], and its antimicrobial activity was evaluated. Phylogenetic and structural analyses revealed that altitudin S is a distinct circular bacteriocin characterized by unique sequence features and physicochemical properties compared to other circular bacteriocins. Homologs of altitudin S were identified across diverse *Bacillales* species, displaying conserved structural features including a five-helix fold, together with comparable physicochemical traits. Their biosynthetic gene clusters also exhibited high conservation, consistently encoding an M48 family metallopeptidase. These findings support the designation of altitudin S as the prototype of a novel subgroup of circular bacteriocins. The discovery and comprehensive characterization of altitudin S expand the known diversity of circular bacteriocins and highlight its potential as a promising candidate for future biotechnological and antimicrobial applications.

## 2. Material and methods

### 2.1. Bacterial isolate and bacteriocin mining

The soil-derived strain *B. altitudinis* ECC22 was originally isolated from soil samples collected by high-school students participating in the Micromundo citizen science project [11,12]. This strain was previously shown to produce the circular bacteriocins pumilarin and altitudin A [9]. To extend these findings, a more comprehensive re-analysis of its genome (GenBank accession number CP137888) was performed using antiSMASH (https://antismash.secondarymetabolites.org/, accessed on 13 January 2025) [13]. The analysis was further supported by BLASTp (NCBI) and UniProt for peptide comparisons, and by SnapGene 6.2.1. (GSL Biotech, San Diego, CA, USA) for the characterization of the identified bacteriocin gene clusters (BGCs).

### 2.2. MALDI-TOF MS and LC-MS/MS analysis of purified supernatants from *B. altitudinis* ECC22

Bacteriocins were purified from the cell-free supernatant (CFS) of *B. altitudinis* ECC22 using a multi-step chromatographic procedure [9]. *B. altitudinis* ECC22 was cultivated for 24 h in brain heart infusion (BHI) broth (Oxoid Ltd., Basingstoke, UK) at 32 °C with agitation at 250 rpm in an orbital shaker (Ecotron, Infors HT, Braunschweig, Germany). The purified CFS fraction obtained from the final RP-FPLC chromatographic step exhibiting antimicrobial activity, was analyzed by MALDI-TOF MS to determine the molecular mass of the peptides. In addition, this fraction was subjected to LC-MS/MS analysis to determine the amino acid sequences of the resulting trypsin-digested peptides at the Unidad de Espectrometría de Masas (CAI Técnicas Biológicas, UCM, Madrid, Spain), as previously described [9].

### 2.3. Cell-free synthesis of altitudin S and evaluation of antimicrobial activity

A synthetic gene construct was designed containing the C-terminal (IC, 36-amino acids) and N-terminal (IN, 104-amino acids) fragments of the natural split Npu DnaE intein flanking the 76-amino acid sequence of mature altitudin S, following a protocol previously described by our research group [10]. In this design, the S39 residue of altitudin S was selected as the +1 residue, while residue I38 was placed in the terminal position, adjacent to the IN fragment. The linear amino acid sequence of the construct was reverse-translated according to the codon usage of *E. coli* using the GeneArt Gene Synthesis tool (Thermo Fisher Scientific). The resulting gene construct was then cloned into a pUC-derived expression vector under the control of a T7 promoter and terminator (Supplementary Fig. S1). The synthetic gene construct cloned in the pUC-derived vector (pCirc-AltS), was obtained from GeneArt (Thermo Fisher Scientific). Plasmid pCirc-AltS served as template for the cell-free synthesis of altitudin S with the PURExpress In Vitro Protein Synthesis Kit (New England Biolabs, Ipswich, MA, USA) [10,14,15]. The DNA template was added at a final concentration of 10 ng/µL in 25 µL reactions, incubated at 37 °C for 2 h and then left at room temperature overnight. The antimicrobial activity of the IV-CFPS/SIML reactions was evaluated using a spot-on-agar test (SOAT) [14]. Briefly, 5 μL of the IV-CFPS/SIML reaction mixture was spotted onto the surface of Petri plates overlaid with a 0.8% soft-agar culture of the indicator microorganism (ca. 10^5^ cfu/mL) (Supplementary Table S1). The plates were then incubated for 24 h, until zones of inhibition were observed.

### 2.4. Bioinformatic analyses of altitudin S and homologs

#### 2.4.1. Identification of homologs

To identify homologs of altitudin S, the mature peptide sequence of this bacteriocin was used as a query in a BLASTp search against the NCBI database. Sequences showing 100% query coverage were considered putative homologs or variants of altitudin S.

#### 2.4.2. Sequence alignment and phylogenetic analysis

To explore the evolutionary relationships of altitudin S and its homologs, the sequences of these peptides, together with all previously described and characterized circular bacteriocins, were aligned using Clustal Omega (EMBL-EBI) with default parameters in the Jalview 2.11.4.1 software [16]. A neighbor-joining phylogenetic tree was generated in Jalview using the BLOSUM62 substitution matrix, and visualized with Interactive Tree of Life (iTOL, https://itol.embl.de).

#### 2.4.3. Physicochemical properties calculations

The physicochemical properties of altitudin S and its homologs were computed using the ProtParam tool available at ExPASy [17]. For each peptide sequence, the following parameters were determined: molecular mass (adjusted by subtracting 18 Da to account for water loss during cyclization), theoretical isoelectric point (pI), net charge at pH 7.0, aliphatic index, and grand average of hydropathicity (GRAVY). These parameters provide insight into peptide stability, hydrophobicity, and electrostatic characteristics, which are key determinants of antimicrobial activity and membrane interaction.

#### 2.4.4. Structural modeling

The three-dimensional (3D) structures of altitudin S and its homologs was predicted using the AlphaFold server [18] and visualized with ChimeraX [19].

#### 2.4.5. Genomic context analysis

The genomes of strains encoding altitudin S homologs were analyzed with antiSMASH, supported by BLASTp (NCBI) and UniProt for peptide comparisons, and SnapGene 6.2.1 for annotation and visualization of the bacteriocin gene clusters (BGCs).

## 3. Results

### 3.1. Genomic features of the altitudin S bacteriocin gene cluster

*B. altitudinis* ECC22 carries a circular chromosome of 3,807,059 bp (GenBank accession number CP137888) and was previously reported to encode three bacteriocin biosynthetic gene clusters (BGCs) corresponding to pumilarin, altitudin A, and a putative closticin 574-like peptide [9]. Among these, pumilarin and altitudin A were experimentally confirmed, whereas the closticin 574-like cluster remained uncharacterized. However, a more detailed re-examination of the genome revealed an additional, previously overlooked BGC predicted to encode a novel circular bacteriocin, which we designated altitudin S.

The predicted precursor of altitudin S is a 132-amino acid peptide encoded within a cluster that also contains genes for ABC transporter proteins, a stage II sporulation protein M (SpoIIM) of the DUF95 superfamily, and a YIP1 family membrane protein (Fig. 1), all characteristic components of circular BGCs. The overall cluster organization closely resembles that of the pumilarin and altitudin A; however, a key distinction is the presence of an additional gene encoding a predicted M48 family metallopeptidase, which is absent from the other two clusters. This unique feature points to a potentially distinct mechanism of leader peptide processing or maturation in altitudin S.

**Fig. 1.**
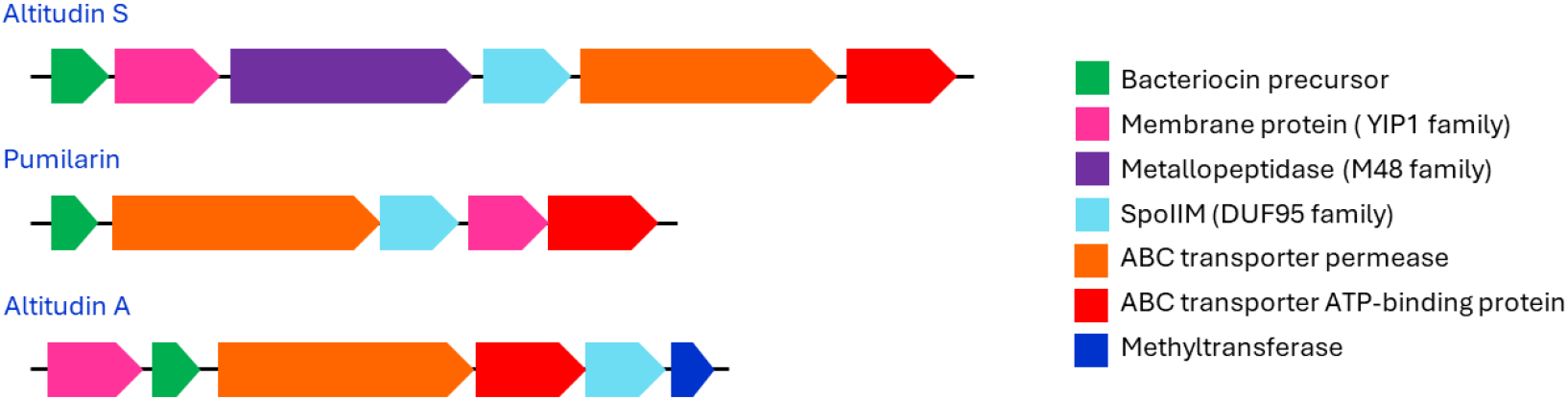
Genomic organization of the altitudin S biosynthetic gene cluster (BGC) for altitudin S compared with pumilarin and altitudin in *B. altitudinis* ECC22. Genes are color-coded by predicted function.

### 3.2. MALDI-TOF MS and LC-MS/MS analyses of purified bacteriocins produced by *B. altitudinis* ECC22

Purification of the cell-free supernatant (CFS) of *B. altitudinis* ECC22 by multi-step chromatography, followed by MALDI-TOF MS analysis of the most active RP-FPLC fraction, revealed three major peptides with molecular masses of 6598.9, 7089.1, and 8381.9 Da, corresponding to altitudin A, pumilarin, and the novel peptide altitudin S, respectively (Fig. 2). For altitudin S, the observed mass was 18 Da lower than the theoretical molecular mass (8397.0 Da), consistent with a dehydration event caused by amide bond formation between the N- and C-termini, resulting in cyclization. Similar mass shifts were observed for altitudin A (6615.9 vs. 6598.9 Da) and pumilarin (7105.4 vs. 7089.1 Da), confirming that all three peptides undergo cyclization in *B. altitudinis* ECC22.

**Fig. 2.**
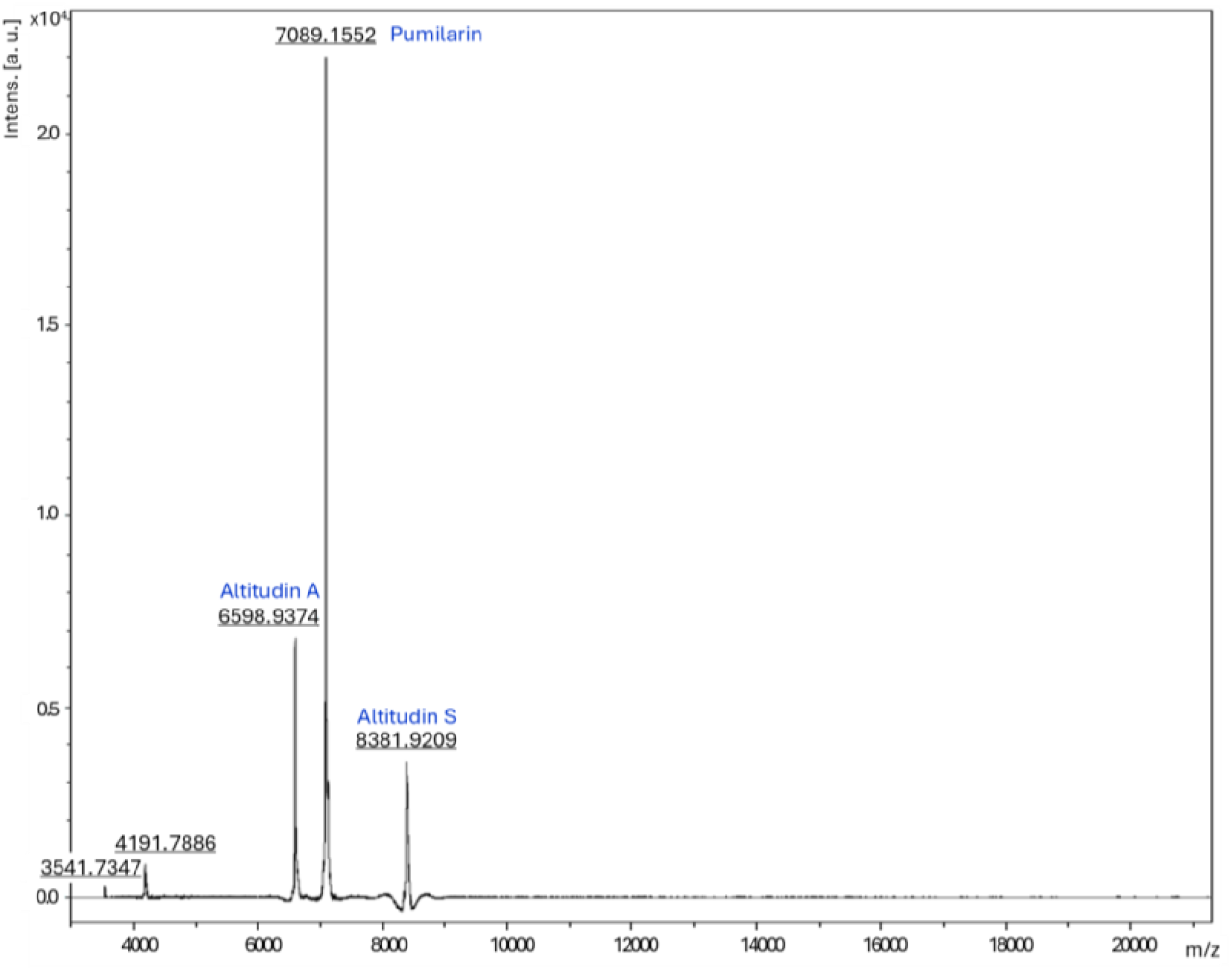
MALDI-TOF MS spectrum of the most active RP-FPLC fraction from the purified cell-free supernatant (CFS) of *B. altitudinis* ECC22, showing peptide peaks corresponding to altitudin A (6598.9 Da), pumilarin (7089.1 Da), and the novel circular bacteriocin altitudin S (8381.9 Da).

Targeted LC–MS/MS analysis of trypsin-digested peptides from the purified CFS further validated the identity of altitudin S. Two peptides, TTWNQAQK and AAVTWLAK, were unambiguously assigned to the predicted mature sequence (Supplementary Table S2). Notably, the detection of AAVTWLAK, which bridges residues L1 and W76, validated the head-to-tail circularization junction, providing definitive evidence of altitudin S production by *B. altitudinis* ECC22. Together, these findings demonstrate that altitudin S is synthesized as a 132-amino acid precursor, which undergoes proteolytic removal of a 56-amino acid leader peptide to yield a 76-amino acid mature circular bacteriocin (Fig. 3A).

**Fig. 3.**
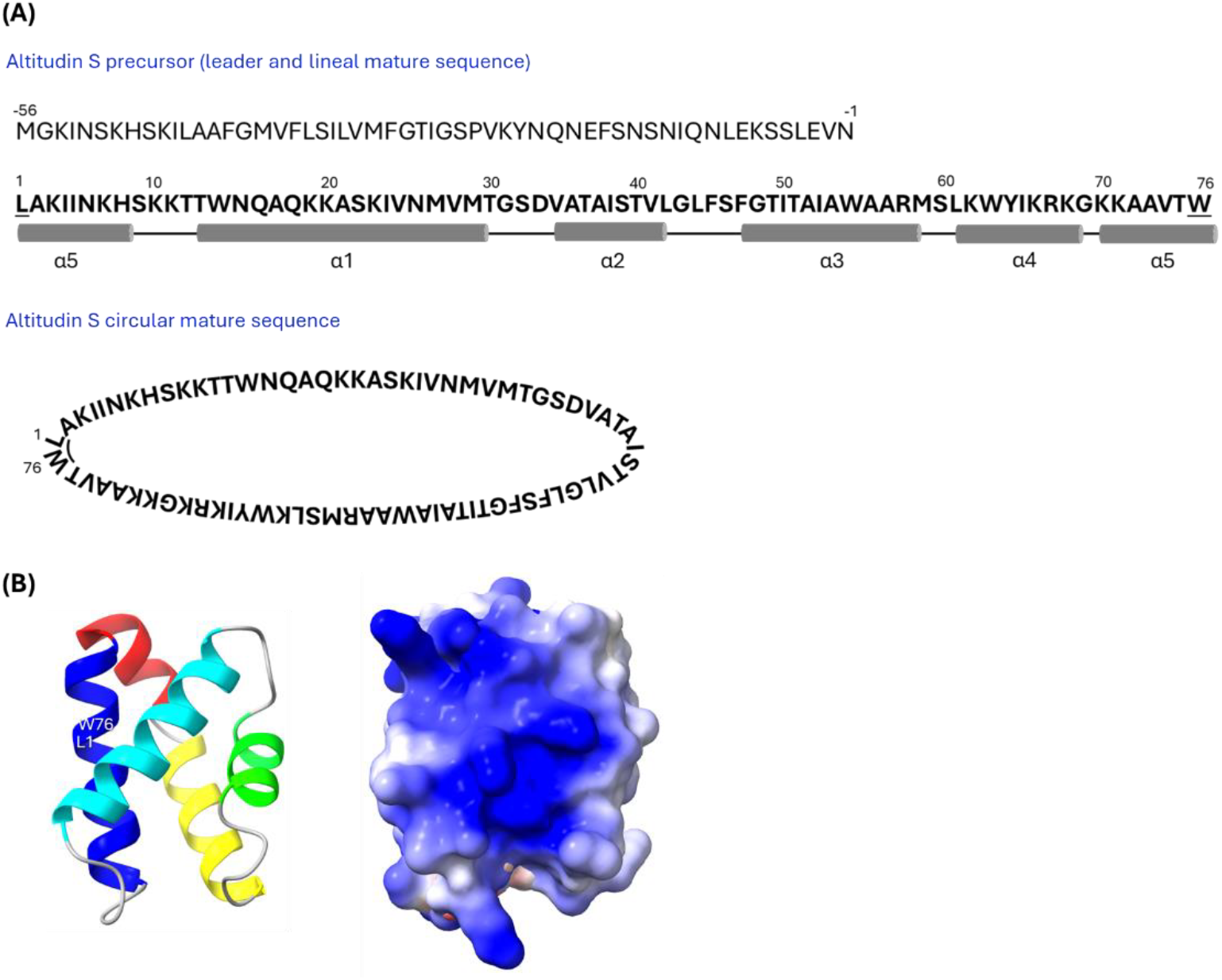
(A) Schematic representation of the altitudin S precursor peptide, composed of a 56-residue leader sequence and a 76-residue mature peptide. The circularization junction (L1-W76) is indicated. Predicted α-helices within the mature sequence are shown in grey. (B) Predicted 3D structure of altitudin S generated with AlphaFold and visualized using ChimeraX. Left: cartoon representation highlighting the five α-helices. Right: electrostatic surface potential, colored red (negative), white (neutral), and blue (positive).

### 3.3. Physicochemical profile and predicted 3D structure of altitudin S

From the ProtParam evaluation of the physicochemical properties of altitudin S, the peptide is predicted to be a highly cationic peptide, with a theoretical pI of 11.0, and a net positive charge of +13, arising from 14 basic residues (2 arginines and 12 lysines) and a single acidic residue. The peptide also exhibits an aliphatic index of 90.0 and a nearly-neutral GRAVY score of 0.025. Collectively, these properties describe a peptide with moderate hydrophobicity but a strongly positive electrostatic profile, physicochemical characteristics commonly associated with antimicrobial activity.

Structural modeling using AlphaFold predicted that altitudin S adopts a saposin-like fold, consisting of five α-helices arranged in a compact circular conformation (Fig. 3B). Electrostatic surface visualization revealed a pronounced positive charge distribution. Visualization of the electrostatic surface highlighted the strongly positive character of the molecule, consistent with its predicted physicochemical properties and likely essential for mediating interactions with negatively charged bacterial membranes.

### 3.4. IV-CFPS/SIML procedure for the synthesis and determination of the antimicrobial activity of altitudin S

Because altitudin S could not be purified to homogeneity from the CFS of *B. altitudinis* ECC22, a synthetic gene construct (pCirc-AltS) was designed and employed as a template for the in vitro cell-free protein synthesis (IV-CFPS) coupled to split intein-mediated ligation (SIML) of the resulting peptide. This approach enabled both the production and head-to-tail circularization of the peptide, thereby facilitating the functional validation of this novel bacteriocin.

The IV-CFPS/SIML-derived altitudin S exhibited antimicrobial activity against a restricted panel of indicator strains, including *Pediococcus damnosus* CECT 4797, *Listeria monocytogenes* CECT 4032, *Paenibacillus larvae* 00223, and *Kocuria rhizophila* CECT 241. In contrast, no inhibitory activity was detected against other Gram-positive bacteria, such as representatives of the genera *Lactococcus, Enterococcus, Staphylococcus, Streptococcus, Bacillus, Erysipelothrix*, and *Clostridium* (Table 1 and Supplementary Fig. S2).

**Table 1.** Antimicrobial activity of IV-CFPS/SIML-produced altitudin S, evaluated using the spot-on-agar test (SOAT) against selected Gram-positive indicator strains.

| Indicator strain | Antimicrobial activity* |
| --- | --- |
| *Pediococcus damnosus* CECT 4797 | + |
| *Lactococcus garvieae* 5806 | - |
| *Enterococcus faecium* ER46 | - |
| *Paenibacillus larvae* DB25 | + |
| *Listeria monocytogenes* CECT 4032 | + |
| *Staphylococcus aureus* ZTA11/00117ST | - |
| *Streptococcus suis* C2969/03 | - |
| *Streptococcus agalactiae* DICM11/00863 | - |
| *Bacillus pumilus* PE12 | - |
| *Bacillus cereus* ICM17/00252 | - |
| *Erysipelothrix rhusiopathiae* ICM21/01900 | - |
| *Kocuria rhizophila* CECT 241 | + |
| *Clostridium perfringens* DICM15/00067-5A | - |
| *Streptococcus mutans* ATCC 25175 | - |

\*Samples showing a clear halo of inhibition (+) or no halo of inhibition (-).

### 3.5. Identification of altitudin S homologs

BLASTp analysis using the mature sequence of altitudin S as a query revealed several homologs across diverse *Bacillales* species, including *B. altitudinis, Bacillus pumilus, Aeribacillus alveayuensis, Evansella* spp., *Siminovitchia sediminis*, and *Caldalkalibacillus mannanilyticus* (Fig. 4). These homologs shared 100% query coverage and ranged from 75 to 82 amino acids (8307–9311 Da). Despite sequence variability, they consistently retained the distinctive physicochemical profile of altitudin S, characterized by highly basic pI values (10.3–11.0), strong cationic charges (+7 to +13), and low or near-neutral hydrophobicity (GRAVY −0.177 to 0.272) (Supplementary Table S3).

**Fig. 4.**
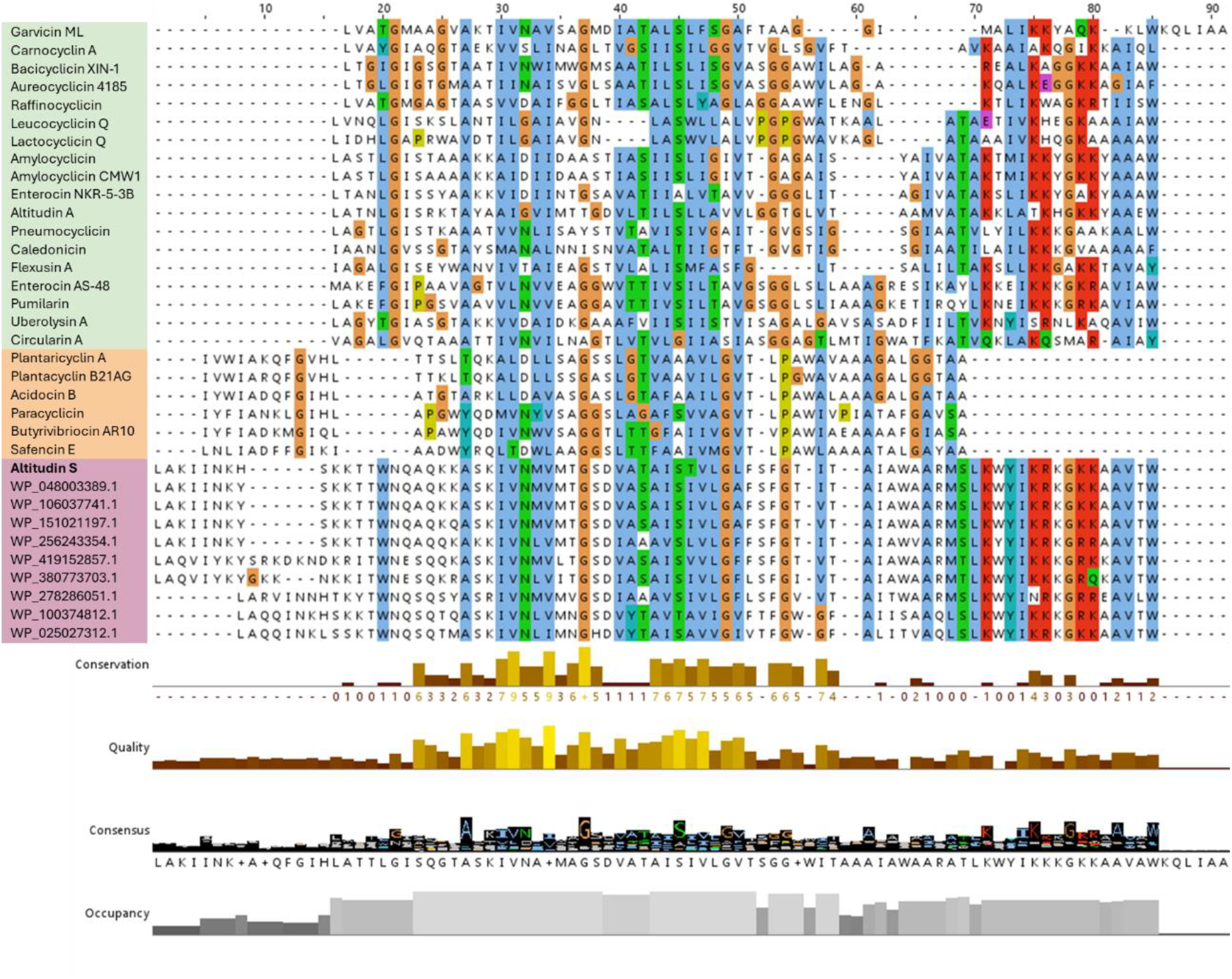
Multiple sequence alignment of altitudin S, its homologs, and representative circular bacteriocins. Mature peptide sequences of subgroup I circular bacteriocins (green box), subgroup II bacteriocins (orange box), and altitudin S with its homologs (purple box) were aligned using the Clustal Omega algorithm implemented in Jalview (v.2.11.4.1) [16]. Amino acid residues are colored according to the Clustal scheme: blue (hydrophobic), green (polar), orange (small hydrophobic), and red (positively charged). Tracks below the alignment provide additional information: Conservation shows residue conservation across all sequences; Quality reflects alignment confidence and physicochemical similarity; Consensus depicts the most frequent amino acid at each position and its conservation as a sequence logo; and Occupancy indicates the proportion of sequences containing a residue (non-gap) at each position.

Predicted three-dimensional (3D) structures of these homologs revealed a conserved compact globular fold composed of five α-helices, consistent with the scaffold identified for altitudin S (Supplementary Fig. S3). This structural conservation indicates that despite some sequence divergence, the tertiary organization remains preserved.

Comparative genomic analysis of the biosynthetic clusters encoding altitudin S and its homologs showed a highly conserved architecture. Each cluster included the structural gene for the precursor peptide, a YIP1 family membrane protein, a SpoIIM (DUF95 superfamily) protein, and ABC transporter subunits (Supplementary Fig. S4). A distinctive hallmark of the altitudin S cluster, shared across all homologs, was the presence of an M48 family metallopeptidase gene located adjacent to the structural gene. This protease is absent from previously characterized circular bacteriocin clusters, such as those encoding pumilarin and altitudin A [9], which otherwise contain conserved components including the ABC transporter and SpoIIM.

### 3.6. Comparative analysis of altitudin S-like homologs with characterized circular bacteriocins

Comparison of the mature amino acid sequences of altitudin S and its homologs with those of all characterized circular bacteriocins revealed limited similarity to either subgroup I (e.g., garvicin ML, enterocin AS-48, circularin A, altitudin A, pumilarin) or subgroup II (e.g., plantaricyclin A, plantacyclin B21AG, acidocin B, paracyclicin). Multiple sequence alignment uncovered unique sequence features that separate altitudin S and its homologs from both canonical subgroups. Notably, these include an extended N-terminal motif (LAKITNK…) and an elevated frequency of lysine residues throughout the sequence, conferring the exceptionally strong cationic character of this group (Fig. 4). Together, these features diverge from the conserved sequence signatures characteristic of subgroup I and II bacteriocins, reinforcing the classification of altitudin S-like peptides as a structurally and functionally distinct lineage of circular bacteriocins.

Phylogenetic analysis of altitudin S, its homologs, and representative circular bacteriocins confirmed their divergence. Altitudin S and its homologs formed a distinct clade that branched separately from the established subgroup I and II clusters (Fig. 5). The extended branch length of this clade reflects substantial evolutionary distance, supporting its recognition as an independent lineage within the circular bacteriocin family.

**Fig. 5.**
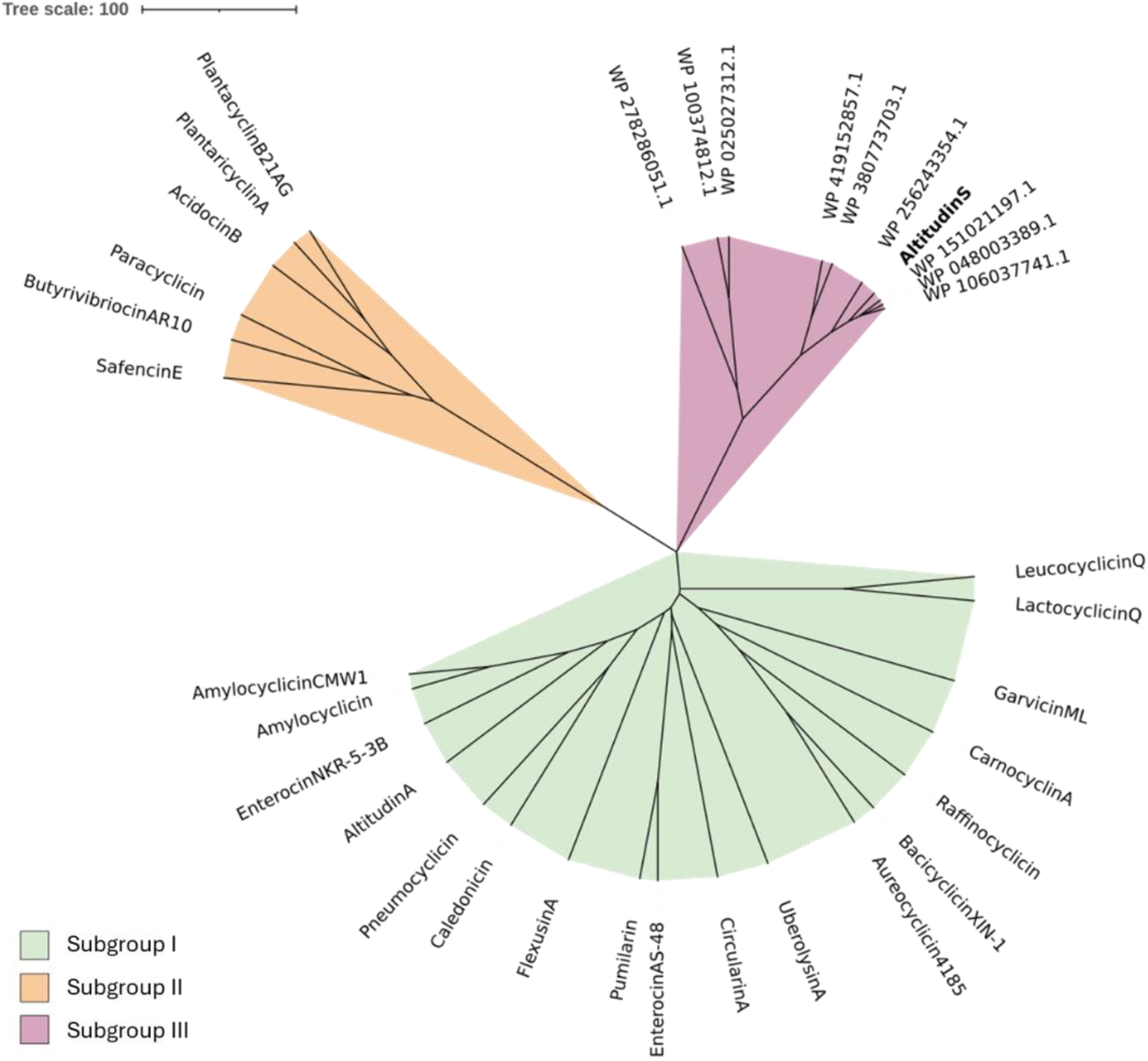
Phylogenetic analysis of circular bacteriocins. Phylogenetic tree derived from the multiple sequence alignment of representative circular bacteriocins and altitudin S homologs. Alignments were generated with Clustal Omega, and the tree was inferred using the Neighbor-Joining algorithm with the BLOSUM62 substitution matrix. Visualization was performed with iTOL (Interactive Tree of Life; https://itol.embl.de). Branch coloring denotes the three proposed subgroups: subgroup I (green), subgroup II (orange), and subgroup III (purple), the latter represented by altitudin S and its homologs.

The physicochemical properties of the altitudin S-like peptides differ markedly from those of known circular bacteriocins. Subgroup I members typically comprise 60–70 amino acids, have highly basic pI values (9.5–10.5), moderate hydrophobicity (GRAVY 0.5–1.1), and carry net positive charges of +2 to +6. Subgroup II members, in contrast are shorter (58 amino acids) and display greater variability in charge and hydropathy, including neutral or negatively charged representatives. Altitudin S-like peptides diverge from both patterns since they are longer (75– 82 amino acids), with extremely high theoretical pI values (10.3–11.0), unusually strong net positive charges observed (+7 to +13), and near-neutral GRAVY scores (Supplementary Table S4), highlighting their unusual basicity and low hydrophobicity.

## 4. Discussion

The global rise in antimicrobial resistance (AMR) poses a significant threat to public health, underscoring the urgent need for new antimicrobial agents. Bacteriocins, especially circular bacteriocins, have garnered considerable attention due to their stability and potential as alternative antimicrobial agents [7]. Here, we report the identification and characterization of a novel circular bacteriocin, altitudin S, produced by *B. altitudinis* ECC22, which was previously reported to produce the circular bacteriocins pumilarin and altitudin A [9], reinforcing *Bacillus* sp. as a valuable source of antimicrobial peptides [20].

A more comprehensive re-analysis of the *B. altitudinis* ECC22 genome revealed a third biosynthetic gene cluster (BGC) predicted to encode a putative new circular bacteriocin, in addition to those for pumilarin and altitudin A. Examination of the altitudin S BGC revealed conserved features typical of circular bacteriocin operons, including genes for the precursor peptide, a SpoIIM protein (DUF95 superfamily), a YIP1 family membrane protein, and a two-component ABC transporter (Fig. 1) [21]. Similar elements are also found in the pumilarin and altitudin A clusters, indicating a shared organizational backbone. However, the altitudin S cluster is distinguished by the presence of a gene encoding an M48 family metallopeptidase adjacent to the structural gene. This protease, absent from pumilarin and altitudin A clusters, points to a possible role in precursor processing and a specialized maturation pathway. Although the mechanisms of leader peptide cleavage and core peptide cyclization remain elusive, further experimental validation is required to clarify the roles of the associated enzymes, as well as additional factors encoded elsewhere in the genome [6,21]. Notably, many class I circular bacteriocin BGCs also harbor an accessory operon of three to four genes encoding a putative ABC transporter complex, typically a permease, an ATP-binding protein, and a predicted extracellular component, as described for enterocin AS-48 (AS-48EFGH). However, functional studies suggest that this complex has no significant impact on bacteriocin production or immunity [8]. The altitudin S gene cluster also lacks a dedicated immunity gene, a feature common to several circular bacteriocin clusters [22,23].

The production of altitudin S by *B. altitudinis* ECC22 was confirmed by purification of the CFS and MALDI-TOF MS analysis of active fractions. Observed molecular masses for altitudin A, pumilarin, and altitudin S were ∼18 Da lower than their deduced values, consistent with dehydration and head-to-tail cyclization (Fig. 2). LC–MS/MS of tryptic fragments identified TTWNQAQK and AAVTWLAK peptides from the predicted mature sequence. Notably, detection of AAVTWLAK, spanning residues L1 and W76, provided direct evidence of head-to-tail circularization of altitudin S in the native producer. These proteomic findings, together with the genome analysis, demonstrate that altitudin S is synthesized as a 132-amino acid precursor with a 56-amino acid leader sequence and a 76-amino acid core peptide. Following leader peptide cleavage and ligation of L1 and W76, the mature bacteriocin is generated (Fig. 3A). Altitudin S is the largest circular bacteriocin described to date, with the mature peptide and leader sequence exceeding the typical ranges reported for circular bacteriocins (58–70 and 2–48 residues, respectively) [8]. Remarkably, altitudin S displays highly unusual physicochemical properties, with a theoretical pI of 11.0 and a net charge of +13, placing it among the most cationic circular bacteriocins reported to date (Supplementary Table S4). Structural predictions further suggests that, like most circular bacteriocins, altitudin S adopts a compact saposin-like fold composed of five α-helices, with the cyclization site buried in a sterically constrained, hydrophobic α-helix (Fig. 3B) [20]. This arrangement likely stabilizes the circular conformation and facilitates membrane disruption through ion leakage, loss of membrane potential, and ultimately cell death [7,20].

Due to the uncertainty of obtaining homogeneous purified altitudin S through reliable multi-chromatographic procedures, an IV-CFPS/SIML procedure was therefore employed for its in vitro production and circularization [9,10,15]. This approach successfully generated active altitudin S, which showed antimicrobial activity against a narrow range of indicator strains (Table 1; Supplementary Fig. S2). Its restricted spectrum was similar to that of altitudin A, in contrast to the broader inhibition of pumilarin [9], likely reflecting differences in charge density and other determinants of bacteriocin-target interactions [6]. These findings also highlight the utility of IV-CFPS/SIML-based systems as a powerful tool for producing and characterizing circular bacteriocins that are difficult to purify from native sources. Nevertheless, a current limitation of this approach is that the concentration of the synthesized peptide remains unknown, and it is possible that the amount obtained was insufficient to fully capture its antimicrobial potential. Further analyses using higher and accurately quantified concentrations will therefore be essential to better define the overall activity and inhibitory spectrum of altitudin S.

The identification of altitudin S homologs across diverse bacterial genera, including *Bacillus, Evansella*, and *Caldalkalibacillus*, suggest that these bacteriocins are more widely distributed than previously anticipated. These homologs share the distinctive properties of altitudin S, including extended mature peptides (75 to 82 amino acids), high pI and strong cationic charge, near-neutral hydrophobicity, a conserved five-helix saposin-like fold, and similar BGC organization (Supplementary Table S2, Fig. S3 and Fig. S4). Taken together, these conserved sequence, structural, and genomic features indicate that altitudin S-like bacteriocins may represent a coherent lineage within the circular bacteriocin family.

Comparative sequence and phylogenetic analyses further reinforced this conclusion. Despite some conserved motifs, altitudin S-like peptides diverged strongly from circular bacteriocins subgroups I and II (Fig 4). In phylogenetic trees, they consistently clustered together but branched separately, giving rise to a long, independent lineage (Fig. 5). The basal branching pattern and substantial evolutionary distance suggest that these peptides originated from an early divergent event or followed an independent evolutionary trajectory relative to other circular bacteriocins [6,8,24].

Moreover, their predicted physicochemical and genomic features further support this separation. Subgroup I bacteriocins are typically moderately hydrophobic and cationic (net charge +2 to +6; pI ∼10), while subgroup II members are shorter (58 amino acids) and exhibit greater variability in charge and hydropathy [6]. By contrast, altitudin S and its homologs exhibit consistently high theoretical pIs (10.3–11.0), strong net positive charges (+7 to +13), and near-neutral GRAVY values, placing them beyond the typical ranges reported for circular bacteriocins [6,8,24]. This profile suggest that classical hydrophobic pore formation may be less favorable, and that their activity instead relies on electrostatic interactions with negatively charged membranes, followed by surface-associated destabilization rather than stable pore insertion. Such a mechanism is consistent with the two-step model proposed for antimicrobial peptides [25], and with evidence that many cationic peptides disrupt membranes via transient leakage and interfacial activity rather than forming stable pores [25–27]. Further biophysical studies will be needed to validate the precise mode of action of altitudin S-like bacteriocins.

Comparative genomic analysis showed that altitudin S and its homologs share a conserved operon structure, including the structural gene, ABC transporter subunits, a SpoIIM protein, and a YIP1 family membrane protein, but are consistently distinguished by the presence of an M48 metallopeptidase adjacent to the precursor gene. Together with their unusually long leader and mature peptides, this feature strongly suggest that altitudin S-like bacteriocins may undergo a specialized maturation pathway distinct from other circular bacteriocins. Since M48 metallopeptidases are associated with intramembrane proteolysis and peptide trimming [21], their consistent presence across homologous clusters points to additional or alternative post-translational processing steps, highlighting new avenues for investigating the biosynthesis and functional diversification of this novel subgroup of circular bacteriocins.

## 5. Conclusions

By combining genomic mining, multi-step purification, mass spectrometry (MALDI-TOF MS, LC–MS/MS), and synthetic biology approaches (IV-CFPS/SIML), we identified and characterized a novel circular bacteriocin, altitudin S, produced by *B. altitudinis* ECC22. Altitudin S is distinguished by its unusual physicochemical profile, clear phylogenetic separation from known circular bacteriocins, and the unique genomic architecture of its biosynthetic cluster. The discovery of closely related homologs across diverse bacterial genera further reinforces the recognition of altitudin S-like peptides as a distinct and previously uncharacterized lineage within the circular bacteriocin family, underscoring their broad taxonomic distribution and likely ecological significance.

## CRediT authorship contribution statement

Conceptualization, E.M.-A., P.E.H. and J.B.; methodology, E.S., N.P., I.L., P.E.H. and J.B.; investigation, E.S., N.P., I.L., E.M.-A. and J.B.; resources, L.M.C., E.M.-A., P.E.H. and J.B.; data curation, E.S., P.E.H. and J.B.; writing—original draft preparation, E.S.; writing—review and editing, P.E.H. and J.B.; supervision, E.M.-A., P.E.H. and J.B.; project administration, J.B. and E.M.-A.; funding acquisition, L.M.C., P.E.H. and J.B. All authors have read and agreed to the published version of the manuscript.

## Ethical approval

This study did not involve human participants or vertebrate animals. Sampling was compliant with the Nagoya Protocol (reference ABSCH-IRCC-ES-270144-1).

## Data availability

Sequence information: The whole genome assembly of *B. altitudinis* ECC22 is deposited in GenBank under the accession number CP137888.

## Acknowledgments

We gratefully acknowledge the Micromundo project from the Universidad Complutense de Madrid and the participating high-school students from IES El Cantizal (Las Rozas, Madrid), under the guidance of their teacher Alicia Cuartero, for their role in the isolation of *B. altitudinis* ECC22. We also thank the project directors, Prof. Víctor Jiménez-Cid and Prof. Jessica Gil-Serna, for coordinating the Micromundo initiative. Thanks also to the Proteomics Unit of the University Complutense of Madrid (UCM), for the technical assistance in the proteomic analysis of samples.

## Appendix A. Supplementary data

Supplementary material associated with this article (Table S1, Table S2, Table S3, Table S4, Fig. S1, Fig. S2, Fig. S3, Fig. S4) can be found, in the online version, at

## Declaration of generative AI and AI-assisted technologies in the manuscript preparation process

During the preparation of this work the authors used ChatGPT (GPT-5, OpenAI) to assist in improving the clarity, grammar, and flow of the text, and to rephrase certain passages for conciseness. After using this tool, the authors carefully reviewed, edited, and validated all content, and take full responsibility for the scientific accuracy and integrity of the published article.

