## Supplementary information for "Altitudin S from *Bacillus altitudinis* ECC22 defines a new subgroup of circular bacteriocins"

(A)

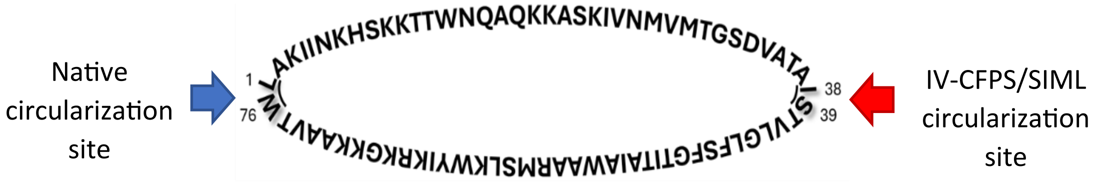


(B)

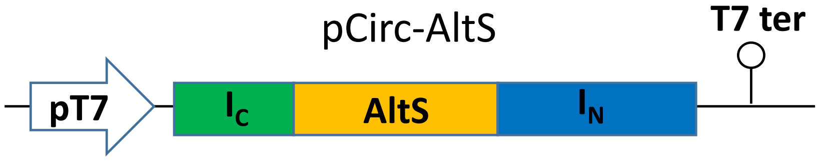


(C)

MIKIATRKYLGKQNVYDIGVERYHNFALKNGFIASNSTVLGLFSFGTITAIAWAARMSLKWYIKRK
GKKAAVTWLAKIINKHSKKTTWNQAQKKASKIVNMVMTGSDVATAICLSYDTEILTVEYGILPIGK
IVEKRIECTVYSVDNNGNIYTQPVAQWHDRGEQEVFEYCLEDGCLIRATKDHKFMTVDGQMMPI
DEIFERELDLMRVDNLPNGT

(D)

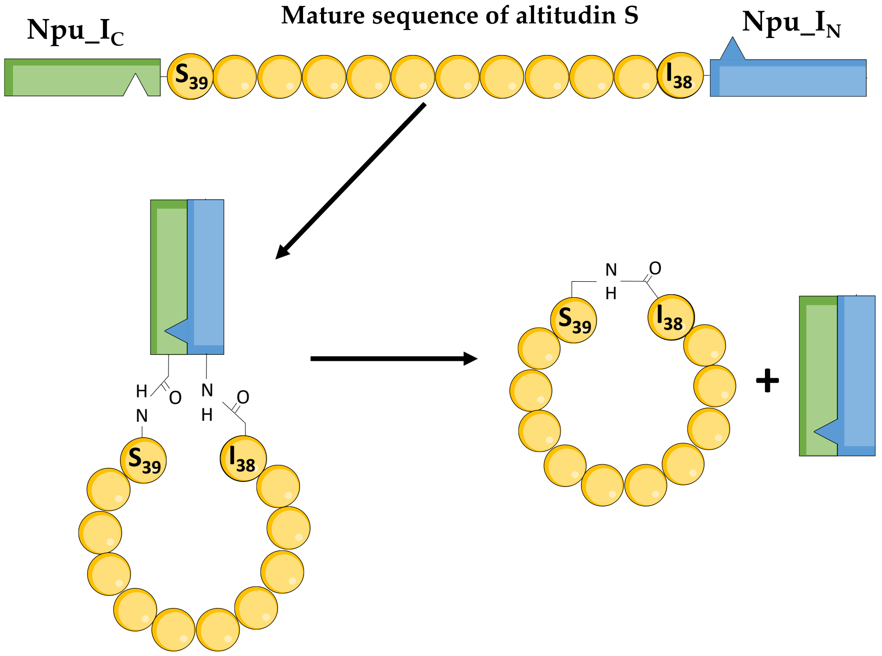


**Supplementary Fig. S1.** Altitudin S (AltS) circularization using the IV-CFPS/SIML system. (A) Amino acid sequence of mature AltS; arrows indicate residues involved in the native head-to-tail circularization (blue arrow), and those selected for IV-CFPS/SIML-mediated production (red arrow). (B) Schematic representation of the pUC-derived protein expression vector (pCirc-AltS) encoding the Npu-AltS fusion construct. (C) Amino acid sequence of the Ic-AltS-$I_N$ construct encoded by pCirc-AltS; the C-terminal 36-amino acids of the Npu split intein (Ic) and the N-terminal 104-amino acids ($I_N$) are shown in green and blue, respectively, with the AltS sequence highlighted in yellow. (D) Schematic overview of the AltS cyclization process. The split intein peptides, $I_N$ and Ic, assemble into an active intein, catalyzing protein splicing and enabling head-to-tail circularization of the mature AltS peptide.

**Supplementary Table S1.** Origin, growth media, and incubation conditions of the bacterial indicator strains used in this study.

| Strain | Origin[a] | Growth medium | Incubation conditions |
|---|---|---|---|
| *Pediococcus damnosus* CECT 4797 | CECT | MRS | 32 °C/ Aerobiosis |
| *Lactococcus garvieae* 5806 | DNBTA | MRS | 32 °C/ Aerobiosis |
| *Enterococcus faecium* ER46 | VISAVET | MRS | 37 °C/ Aerobiosis |
| *Paenibacillus larvae* DB25 | DNBTA | BHI | 37 °C/ Aerobiosis |
| *Listeria monocytogenes* CECT 4032 | CECT | BHI | 37 °C/ Aerobiosis |
| *Staphylococcus aureus* ZTA11/00117ST | VISAVET | BHI | 37 °C/ Aerobiosis |
| *Streptococcus suis* C2969/03 | VISAVET | BHI | 37 °C/ Aerobiosis |
| *Streptococcus agalactiae* DICM11/00863 | VISAVET | BHI | 37 °C/ Aerobiosis |
| *Bacillus pumilus* PE12 | DNBTA | BHI | 37 °C/ Aerobiosis |
| *Bacillus cereus* ICM17/00252 | VISAVET | BHI | 37 °C/ Aerobiosis |
| *Erysipelothrix rhusiopathiae* ICM21/01900 | VISAVET | BHI | 37 °C/ Aerobiosis |
| *Kocuria rhizophila* CECT 241 | CECT | BHI | 37 °C/ Aerobiosis |
| *Clostridium perfringens* DICM15/00067-5A | VISAVET | BHI | 37 °C/ Anaerobiosis |
| *Streptococcus mutans* ATCC 25175 | ATCC | BHI | 37 °C/ Anaerobiosis |

[a] DNBTA: Departamento de Nutrición, Bromatología y Tecnología de los Alimentos, Facultad de Veterinaria, Universidad Complutense de Madrid (UCM), Madrid, (Spain). VISAVET: Centro de Vigilancia Sanitaria Veterinaria, Universidad Complutense de Madrid (UCM), Madrid, (Spain). CECT: Colección Española de Cultivos Tipo, Valencia, (Spain). ATCC: American Type Culture Collection.

**Supplementary Table S2.** LC–MS/MS identification of altitudin S-derived peptides. Tryptic peptides detected in the purified CFS of *B. altitudinis* ECC22 were analyzed by LC–MS/MS. Two peptides, TTWNQAQK and AAVTWLAK, were identified, both mapping to the predicted mature sequence of altitudin S (in yellow). Notably, the peptide AAVTWLAK spans residues L1 and W76, thereby confirming the head-to-tail circularization junction of the bacteriocin. Theoretical MH$^+$ values and corresponding detected m/z are reported.

LAKIINKHSKKTTWNQAQKKASKIVNMVMTGSDVATAISTVLGLFSFGTITAIAWAARMSLKWYIKRKGKKAAVTW

| Peptide Sequence | Theoretical MH+ (Da) | Detected m/z |
|---|---|---|
| AAVTWLAK | 859.50361 | 430.25534 |
| TTWNQAQK | 976.48467 | 488.74573 |

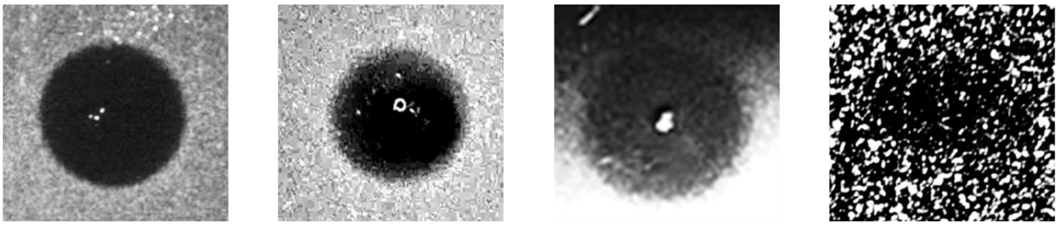

*P. damnosus* CECT 4797 *L. monocytogenes* CECT 4032 *P. larvae* DB25 *K. rhizophila* CECT 241

**Supplementary Fig. S2.** Antimicrobial activity of the IV-CFPS/SIML-produced altitudin S by using the spot-on-agar test (SOAT) against the indicator strains *P. damnosus* CECT 4797, *L. monocytogenes* CECT 4032, *P. larvae* DB25, and *K. rhizophila* CECT 241. Zones of growth inhibition surrounding the application spots indicate antimicrobial activity.

**Supplementary Table S3.** General properties of altitudin S homologs identified by BLASTp. Amino acid sequences and predicted physicochemical parameters of altitudin S and its homologs identified across diverse *Bacillales* species. Reported properties include mature peptide length (aa), molecular mass (Da; corrected for circularization by subtracting 18 Da), theoretical isoelectric point (pI), net charge at pH 7.0, aliphatic index, and GRAVY (Grand Average of Hydropathy). Homologs were retrieved from GenBank, and their corresponding accession numbers and producer organisms are indicated.

| **Bacteriocin** | **Producer organism** | **Mature sequence** | **Length of mature sequence (aa)** | **Molecular mass (Da)** | **Theore-tical pI** | **Net charge** | **Aliphatic index** | **GRAVY** | **Genome GenBank** |
|---|---|---|---|---|---|---|---|---|---|
| Altitudin S | *Bacillus altitudinis* | LAKIINKHSKKTTWNQAQKKASKIVNMVMTGSDVATAISTVLGLFSFGTITAIAWAARMSLKWYIKRKGKKAAVTW | 76 | 8379 | 11.00 | 13 | 90.00 | 0.025 | CP137888 |
| WP_048003389.1 | *Bacillus altitudinis* | LAKIINKYSKKTTWNQAQKKASKIVNMVMTGSDVASAISIVLGAFSFGTITAIAWAARMSLKWYIKRKGKKAAVTW | 76 | 8361 | 10.75 | 13 | 91.32 | 0.091 | JAWVNE010000001 |
| WP_106037741.1 | *Bacillus pumilus* | LAKIINKYSKKTTWNQAQKKASKIVNMVMTGSDVASAISIVLGAFSFGTVTAIAWAARMSLKWYIKRKGKKAAVTW | 76 | 8347 | 10.75 | 13 | 90.00 | 0.087 | VKQA01000100 |
| WP_151021197.1 | *Bacillus pumilus* | LAKIINKYSKKTTWNQAQKQASKIVNMVMTGSDVASAISIVLGAFSFGTVTAIAWAARMSLKWYIKRKGKKAAVTW | 76 | 8347 | 10.72 | 12 | 90.00 | 0.092 | JAKOBC010000001 |
| WP_256243354.1 | *Bacillus* spp. | LAKIINKYSKKTTWNQAQQKAKKIVNLVMTGSDIAAAVSLVLGFFSFGTITAIAWVARMSLKYYIKRKGRRAAVTW | 76 | 8505 | 10.95 | 13 | 98.95 | 0.129 | JAETXM010000001 |
| WP_419152857.1 | *Aeribacillus alveayuensis* | LAQVIYKYSRKDKNDKRITWNESQQKASKIVNMVLTGSDVASAISVVLGFFSFGTVTAIAWAARMTLKWYIKKKGRKKAVTW | 82 | 9311 | 10.42 | 12 | 86.83 | -0.177 | JAUSTR010000001 |
| WP_380773703.1 | *Siminovitchia sediminis* | LAQVIYKYGKKNKKITWNESQKRASKIVNLVITGSDIASAISIVLGFLSFGIVTAIAWAARMTLKWYIKKKGRQKAVTW | 79 | 8883 | 10.58 | 13 | 107.47 | 0.086 | JBHUEO010000001 |
| WP_278286051.1 | *Evansella* spp. | LARVINNHTKYTWNQSQSYASRIVNMVMVGSDIAAAVSIVLGFLSFGVVTAITWAARMSLKWYINRKGRREAVLW | 75 | 8471 | 11.00 | 7 | 104.00 | 0.272 | JARUKY010000001 |
| WP_100374812.1 | *Bacillus* spp. | LAQQINKHSKKTWNQSQTQASKIVNLVMNGSDVYTAVTAVIGVFTFGWGFAIISAAQLSLKWYIKKKGRKAAVTW | 75 | 8311 | 10.42 | 9 | 91.07 | -0.005 | KZ454938 |
| WP_025027312.1 | *Caldalkalibacillus mannanilyticus* | LAQQINKLSSKTWNQSQTMASKIVNLIMNGHDVYTAISAVVGIVTFGWGFALITVAQLSLKWYIKRKGKKAAVTW | 75 | 8307 | 10.36 | 8 | 105.33 | 0.219 | BAMO01000108 |

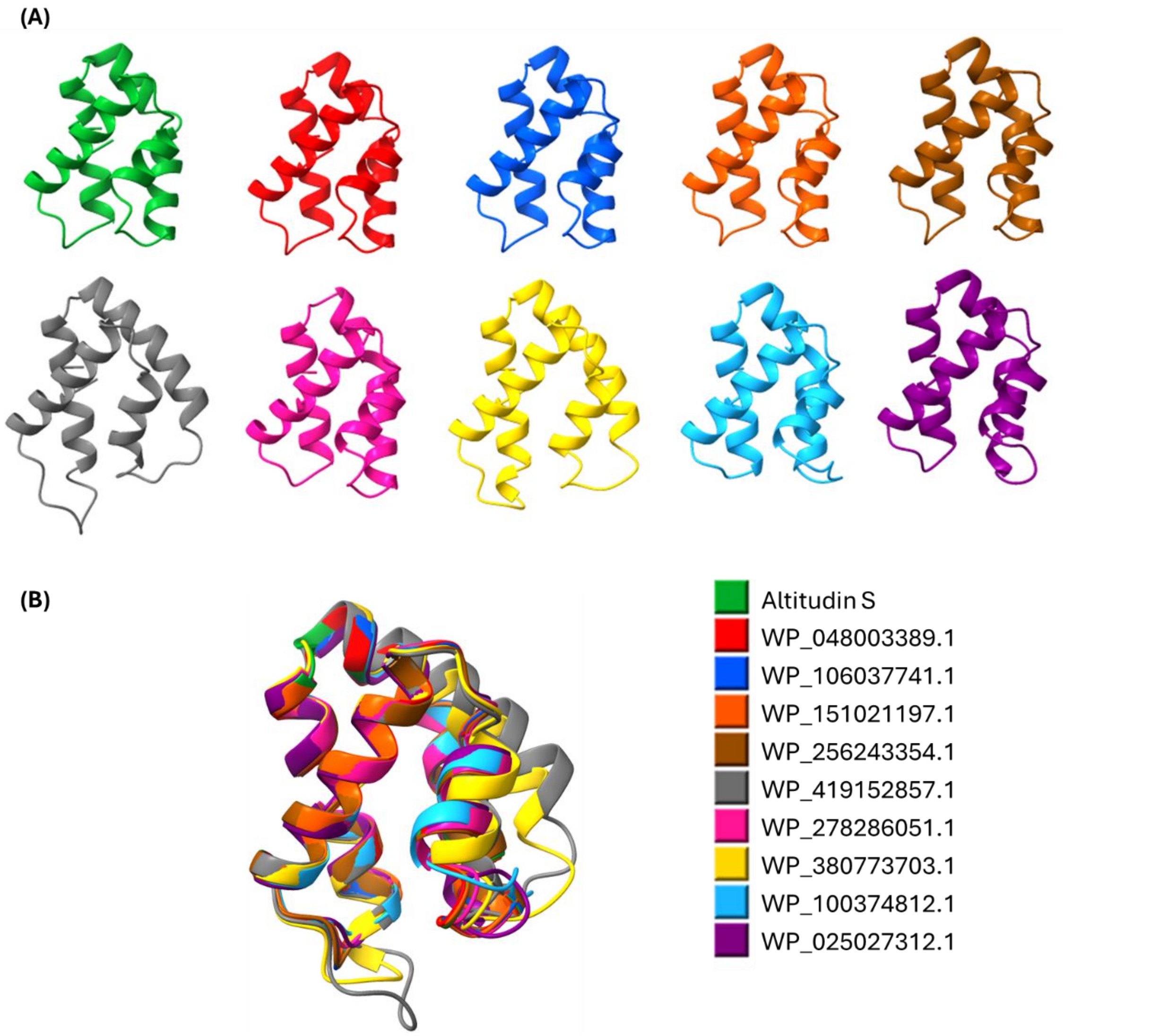


**Supplementary Fig. S3.** Structural analysis of altitudin S and putative homologs (WP_048003389.1, WP_106037741.1, WP_151021197.1, WP_256243354.1, WP_419152857.1, WP_278286051.1, WP_380773703.1, WP_100374812.1, WP_025027312.1). **(A)** Predicted 3D structures of altitudin S and homologous circular bacteriocins, generated using the AlphaFold Server and visualized in ChimeraX. **(B)** Structural superposition of the peptides, aligned and rendered in ChimeraX. Each peptide is indicated by a distinct color.

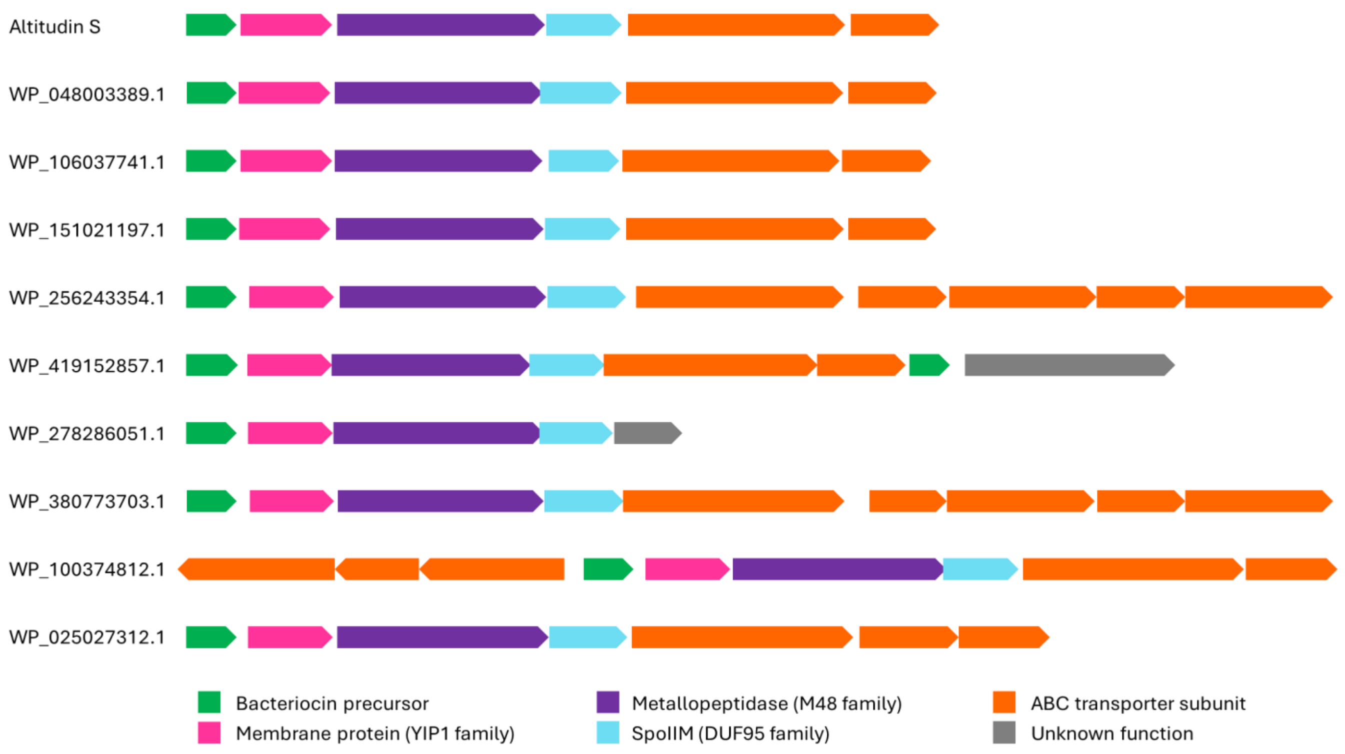


**Supplementary Fig. S4.** Gene clusters for altitudin S and putative homologs (WP_048003389.1, WP_106037741.1, WP_151021197.1, WP_256243354.1, WP_419152857.1, WP_278286051.1, WP_380773703.1, WP_100374812.1, WP_025027312.1). Open reading frames (ORFs) are depicted as arrows, with color coding according to their predicted functions.

**Supplementary Table S4.** General physicochemical properties of circular bacteriocins. Reported producer organisms and corresponding GenBank accession numbers are indicated.

| Bacteriocin | Sub-group | Length of leader sequence (aa) | Length of mature sequence (aa) | Molecular mass (Da) | Theoretical pI | Net charge | Aliphatic index | GRAVY | Producer organism/ strain | Isolation source[a] | Reference |
|---|---|---|---|---|---|---|---|---|---|---|---|
| Altitudin S | | 56 | 76 | 8379 | 11.0 | 13 | 90.0 | 0.025 | *Bacillus altitudinis* ECC22 | Soil | Present study |
| Garvicin ML | i | 3 | 60 | 6007 | 10.1 | 5 | 115.8 | 0.887 | *Lactococcus garvieae* DCC43 | Duck intestines | [1] |
| Carnocyclin A | i | 4 | 60 | 5862 | 10.0 | 4 | 144.7 | 1.058 | *Carnobacterium maltaromaticum* UAL307 | Fresh pork | [2] |
| Bacicyclicin XIN-1 | i | 4 | 60 | 5852 | 10.3 | 3 | 115.8 | 0.877 | *Bacillus* sp. Xin1 | Soil | [3] |
| Aureocyclicin 4185 | i | 4 | 60 | 5608 | 10.0 | 3 | 125.5 | 0.973 | *Staphylococcus aureus* 4185 | Bovine mastitis | [4] |
| Raffinocyclicin | i | 3 | 61 | 6092 | 9.5 | 2 | 117.1 | 0.872 | *Lactococcus raffinolactis* APC 3967 | Raw milk | [5] |
| Leucocyclicin Q | i | 2 | 61 | 6115 | 9.5 | 2 | 131.5 | 0.744 | *Leuconostoc mesenteroides* TK41401 | Japanese pickles | [6] |
| Lactocyclicin Q | i | 2 | 61 | 6060 | 9.7 | 2 | 126.7 | 0.826 | *Lactococcus* sp. QU 12 | Cheese | [7] |
| Amylocyclicin | i | 48 | 64 | 6382 | 9.8 | 5 | 125.5 | 0.850 | *Bacillus amyloliquefaciens* FZB42 | Soil | [8] |
| Amylocyclicin CMW1 | i | 47 | 64 | 6352 | 9.8 | 5 | 127.0 | 0.889 | *Bacillus amyloliquefaciens* CMW1 | Japanese fermented soybean paste | [9] |
| Enterocin NKR-5-3B | i | 23 | 64 | 6317 | 9.9 | 5 | 134.4 | 0.953 | *Enterococcus faecium* NKR-5-3B | Thai fermented fish | [10] |
| Altitudin A | i | 49 | 65 | 6598 | 10.0 | 5 | 123.2 | 0.723 | *Bacillus altitudinis* ECC22 | Soil | [11] |
| Pneumocyclicin | i | 34 | 64 | 6156 | 10.1 | 5 | 137.3 | 1.128 | *Streptococcus pneumoniae* (various) | ND | [12] |
| Caledonicin | i | 32 | 64 | 6077 | 10 | 3 | 120.8 | 0.995 | *Staphylococcus caledonicus* APC 4137 | Healthy domestic dog | [13] |
| Flexusin A | i | 6 | 60 | 6098 | 9.6 | 3 | 130.3 | 0.992 | *Bacillus flexus* R29-2 | Marine environment | [14] |
| Enterocin AS-48 | i | 35 | 70 | 7150 | 10.1 | 6 | 117.1 | 0.539 | *Enterococcus faecalis subsp. liquefaciens* S-48 | Human wound exudate | [15] |
| Pumilarin | i | 38 | 70 | 7087 | 10.0 | 5 | 125.4 | 0.579 | *Bacillus pumilus* B4107 | Reisentopf with chicken | [16] |
| Uberolysin A | i | 6 | 70 | 7048 | 9.6 | 3 | 132.6 | 0.937 | *Streptococcus uberis* 42 | Cow mammary secretion | [17] |
| Circularin A | i | 3 | 69 | 6771 | 10.5 | 4 | 124.6 | 1.007 | *Clostridium beijerinckii* ATCC 25752 | Soil | [18] |
| Plantaricyclin A | ii | 33 | 58 | 5571 | 8.6 | 1 | 128.1 | 1.057 | *Lactobacillus plantarum* NI326 | Olives | [19] |
| Plantacyclin B21AG | ii | 33 | 58 | 5668 | 10.0 | 2 | 131.4 | 1.002 | *Lactobacillus plantarum* B21 | Vietnamese fermented sausage | [20] |
| Acidocin B | ii | 33 | 58 | 5622 | 6.8 | 0 | 121.7 | 1.036 | *Lactobacillus acidophilus* M46 | Human feces | [21] |
| Paracyclicin | ii | 24 | 58 | 5907 | 6.7 | 0 | 114.5 | 1.003 | *Lactobacillus paracasei* subsp. *paracasei* DSM 5622 | ND | [22] |
| Butyrivibriocin AR10 | ii | 22 | 58 | 5982 | 4.0 | -2 | 114.7 | 1.002 | *Butyrivibrio fibrisolvens* AR10 | Rumen | [23] |
| Safencin E | ii | 35 | 58 | 5998 | 4.4 | -1 | 118.3 | 0.979 | *Bacillus safensis* APC 4099 | Bees' gut | [24] |

[a]ND: no data.
